# OpenCafeMol: A coarse-grained biomolecular simulator on GPU with its application to vesicle fusion

**DOI:** 10.1101/2025.02.20.639390

**Authors:** Yutaka Murata, Toru Niina, Shoji Takada

## Abstract

There has been an increasing demand for longer-timescale molecular dynamics (MD) simulations of larger biomolecular systems. To meet these demands, using the C++ API of OpenMM, we developed a fast and flexible MD software, OpenCafeMol, for residue-resolution protein and lipid models that shows high performance on graphics processing unit (GPU) machines. We validated OpenCafeMol for folding small proteins, lipid membrane dynamics, and membrane protein structures. Benchmark tests of the computation times showed that OpenCafeMol with one GPU for proteins and lipid membranes can be approximately 100 and 240 times faster than the corresponding simulations on a typical CPU machine (eight cores), respectively. Taking advantage of the high speed of OpenCafeMol, we applied it to two sets of vesicle fusion simulations; one driven by force and the other coupled with conformational dynamics of a SNARE complex. In the latter, a direct MD simulation at a high temperature resulted in vesicle docking, pore formation followed by fusion, which are coupled with local folding of linkers in the SNARE complex. This opens up new avenue to study membrane-fusion mechanisms via MD simulations. The source code for OpenCafeMol is fully available.

## INTRODUCTION

In recent years, there has been an increasing demand for longer-timescale molecular dynamics (MD) simulations of larger systems, especially in molecular biology. Rather than a single protein in aqueous solution, large protein complexes, crowded macromolecular systems, and biomolecular condensates have become the target of MD simulations. To meet these demands, two major approaches have been adopted: 1) speeding up computation using new computers and algorithms, and 2) coarse-graining the model representation. For the former, the special-purpose computer ANTON (1), general-purpose supercomputers (see, for example (2)), and graphics processing unit (GPU) (3, 4) have been successfully employed. For the latter, numerous coarse-grained models with different resolutions have been developed (5–16), along with methodologies to bridge different resolutions (17–21). There is no single universal coarse-grained model for biomolecules; however, different models with different resolutions are useful for different target systems.

Despite intensive studies on each of the two approaches, the combination of the two approaches has rarely been studied. Recently, GPU-accelerated coarse-grained (CG) MD simulations have been realized, showing promise over conventional CPU-based simulations. For this purpose, OpenMM provides a fast, flexible, and multiplatform framework to achieve such development (OpenMM 7 (22)). For example, OpenAWSEM and Open3SPN use the OpenMM Python library and show marked acceleration in CG protein and DNA simulations, respectively (23). OpenABC further extends this direction to extend its applicability (24).

In this study, along the same lines, we developed a fast and flexible MD simulator, called OpenCafeMol, for residue-resolution coarse-grained models of proteins and lipids using the OpenMM library. OpenCafeMol implements a basic set of models used in the previously developed CafeMol software, such as AICG2+, iSoLF (25), and the HPS model (12, 26). For a few test systems of proteins, lipid membranes, and their mixtures, we performed MD simulations with OpenCafeMol, compared the results with those of CafeMol (5), and compared the computation time of OpenCafeMol with that of CPU-based software using typical laboratory-owned computers. We then applied OpenCafeMol to two sets of vesicle fusion simulations; one is the vesicle fusion by an external force and the other is that induced by a SNARE complex. In the latter, a disorder-to-order transition in a linker region of a SNARE complex drives the vesicle pore formation inducing vesicle fusion at a high temperature, but not at a physiological temperature. Without the use of GPU, this type of simulations would not been easily performed. The source code of the OpenCafeMol software is fully available on GitHub.

## METHODS

### The protein model AICG2+

The force field used for the protein benchmark was the AICG2+ model and the Debye–Hückel electrostatic interaction(11). The AICG2+ is a structure-based CG model where one amino acid is represented by one CG particle located on its Cα atom. This can be expressed using the following function:

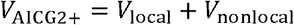

where *V*_local_ represents the potential related to neighboring particles connected by covalent bonds and can be written as follows:

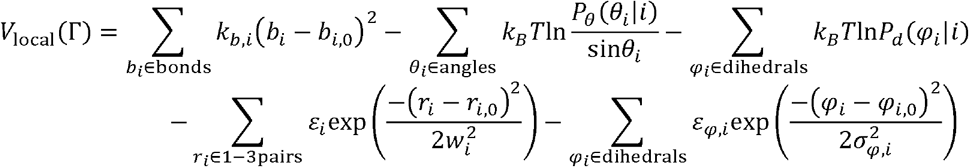

Here, Γ collectively represents the coordinates of all the particles in the target protein. The first term represents the harmonic potential required to limit the virtual bond lengths. The second and third terms are statistical potentials for virtual bond angles and virtual dihedral angles, respectively, representing amino acid sequence-dependent properties, where *P*_*θ*_ (*i*) and *P*_*d*_ *(i)* are probability distributions of angles and dihedral angles that depend on local amino acid sequences. The fourth and fifth terms represent native structure-specific local interactions for the so-called 1-3 pairs and dihedral angles, respectively. *V*_nonlocal_ is the potential for nonlocal particles along the chain and is represented as

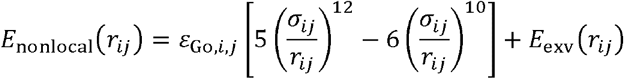

The first term is structure-based Gō-like potential, where *r*_*ij*_ is the distance between residues that are interacting with each other in the native structure. *σ*_*ij*_ is the corresponding distance in the native structure. *ε*_Go,*i*,*j*_ is the energy coefficient that depends on atomic interaction between two residues at the native structure. The second term is an excluded volume potential and can be written as follows:

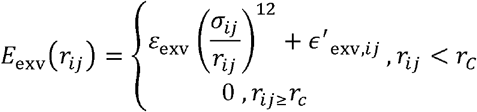

*σ*_*ij*_ is excluded volume distance and depends on residue type. *ε*_exv_ is force coefficient and 0.6 kcal/mol, *r*_*C*_is the cutoff distance and corresponds to 2 *σ*_*ij*_, *ϵ*^′^_exv,*ij*_ is a correction for the cutoff and corresponds to (1/2)^12^*ε*_exv_.

The Debye–Hückel electrostatic interaction can be written as following function.

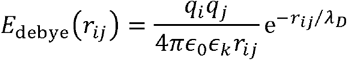

*r*_*ij*_ is the distance between two non-bonded charged particles and *ϵ*_0_ is the dielectric permittivity of vacuum. *ϵ*_*k*_ is the relative permittivity of the solution. *λ*_*D*_is the Debye length and is defined by 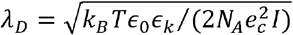, where *N*_*A*_ is the Avogadro’s number, *e*_*c*_ is the elementary charge, and *I* is the ionic strength of the solution.

For the protein simulation, we used the Langevin integrator in OpenMM and 0.01 cafetime^-1^ for the friction coefficient of all particles. The default integrator and friction coefficients were used in GENESIS and CafeMol. We used 0.1 cafetime as the MD step size Δt with the TOML input interface.

### The membrane model iSoLF

For the simulation including the membrane, we used the iSoLF force field for lipids(25). The iSoLF force field can be expressed as follows:

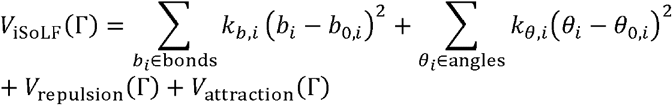

The first and second terms are the harmonic potentials to restrain the virtual bond lengths and the harmonic potential for the virtual bond angles, respectively. *V*_repulsion_ is the Weeks– Chandler Andersen (WCA) potential as a short-range repulsion; explicitly, it has the formula:

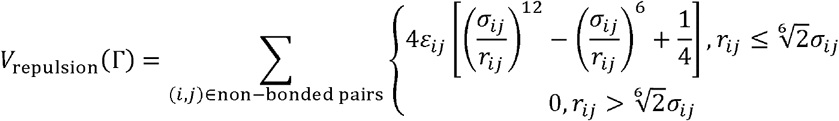

where *σ*_*ij*_ is the arithmetic mean (*σ*_*i*_ + *σ*_*j*_)/2 and *ε*_*ij*_ is the geometric mean 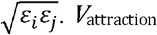 is a short-range attraction potential and can be expressed as follows:

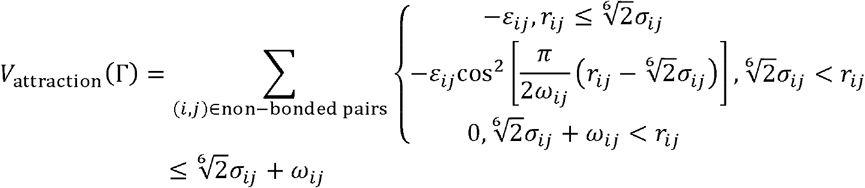

where *ω*_*ij*_ is the range over which the particles are subject to attractive force.

The integrator in the CafeMol simulation for the iSoLF is the method developed by Gao et al. (27). The barostat was semi-isotropic pressure coupling for XY direction and Z was fixed, called Nγ_xy_L_z_T ensemble. For OpenCafeMol, we used the same integrator as that of the protein, and the friction coefficient was 0.005 cafetime^-1^ for all particles. We used the periodic boundary box and MonteCarloMembraneBarostat with XYIsotropic, Z fixed. For the lipid-only simulation, Δt is 0.1 cafetime. In initial configuration, all the lipids are arranged in a square pattern with equal spacing and the box size was tuned so that the APL is 64 □^2^/lipid.

To analyze the tilt angle of rhodopsin in the membrane, we defined the tilt angle as the angle between the z-axis and the line between the rhodopsin N-terminal- and C-terminal particles.

### Protein-lipid interactions

For the protein and lipid simulations, we used the lipid-protein interaction force field developed by Ugarte et al. (28). The force field is expressed as follows:

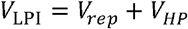

*V*_*rep*_ is the repulsive term and modeled by WCA potential in the same way as iSoLF forcefield, except for 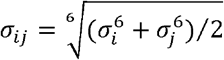 is the hydrophobic-hydrophilic potential and can be expressed as follows:

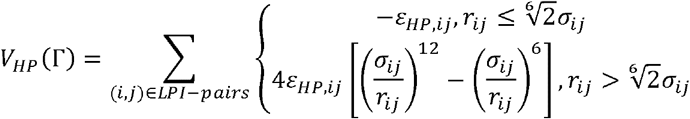

*σ*_*ij*_ is the same as of *V*_*rep*_ . *ε* _*HP*,*ij*_ is specific value for the particle pairs. In this model, each lipid was composed of five particles and 20 types of natural amino acids for a total of 100 parameters. We used 0.2 cafetime as Δt. The initial conditions were prepared using the following steps. First, 128 lipid molecules were prepared in the same manner as for the membrane benchmark. Rhodopsin was then placed on the membrane, and the overlapping lipids were removed. This system was treated as one unit and aligned with 4, 9, 16, and 25 units in the shape of a square. Before the production run, the initial configuration was equilibrated with 1000 steps under NVT conditions.

### HPS model

The HPS model(26) was implemented for simulation of interacting disordered regions in OpenCafeMol. The potential function for this model is expressed as follows:

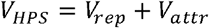

*V*_*rep*_ is the repulsive term and modeled by WCA potential in the same way as iSoLF forcefield, and *V*_*attr*_ is hydrophobic-hydrophilic potential and can be modeled by the same formulation of *V*_*HP*_ in the protein-lipid interaction.

### Simulations of vesicle fusion by force

For the vesicle simulation, we first built a vesicle configuration by aligning lipid bilayer on the sphere, of which the inner radius is 50 □ with 64 □^2^ area per lipid. Then, we placed two vesicles 250 □ apart along x axis. We put a large pseudo sphere at the center of each vesicle with the WCA repulsive force to lipid particles; the parameter of WCA potential is *σ*_pseudo_ = 40 □ *ε* _pseudo_ = 10.0 kcal/mol. To pull one vesicle toward the other, we applied a spring force to the two pseudo spheres as follows:

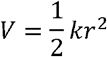

where *r* is the distance between two spheres, and *k* = 0.01 kcal/mol/□^2^ is the spring constant for the pulling.

For the analysis of the center of mass of lipids belongs to each vesicle, first, we divide lipids into two groups that belong to two vesicles by the x coordinates in the initial configuration. After that, geometric center of mass of each group was calculated. To avoid the error from few lipid molecules that went out the vesicle, lipid molecules that contain CG particles >120□Å away from the corresponding geometric center were excluded from the analysis. We monitored the distance between geometric centers of mass of the two groups.

### Simulations of vesicle fusion with SNARE complex

For simulation setup of the vesicle with SNARE complex, we began with lipids configuration prepared in the above section; simulations of vesicle fusion by force. Then, we modeled SNARE complex structure from cis conformation of SNARE complex structure (PDBID: 3HD7) (29). In this modelling, we used MODELLER for building missing atoms and the flexible linker region between two *α*-helix in SNAP-25 which form a helix bundle (30). Next, from this model, we made C *α* model of SNARE complex and the parameter set for AICG2+ forcefield. Some part of sequence was considered as intrinsically disordered region (IDR), 92-94 amino-acid for Synaptobrevin, 255-265 amino-acid for Syntaxin-1A, 82-139 amino-acid for SNAP-25 in the UniProt indexing scheme (31). After this, we set the model of SNARE complex and pulled each terminal of transmembrane helix of Synaptobrevin and Syntaxin-1A to different vesicle by pulling force with the simulation using force field generated from previous part. In this simulation, the force field for the other part, for example lipid-protein interaction, was the same as that in the previous simulations. For the electrostatic interactions, we added charge to residues depending on the type (+1 for Arg and Lys, and -1 for Asp and Glu).

For the production runs, we set partial charges to amino acids in the globular domains of the SNARE complex using the RESPAC method: For each globular domain of Synaptobrevin, Synataxin-1A and SNAP-25, we parametrized charges to fit the electro-static potential of each all-atom structure(32). For the IDR part, we used integer charge, the same as that in the pulling simulation. With this setup, we conducted production runs with 10^8^ MD steps; two replicates in each of temperature, 303 K and 363 K.

## RESULTS

### OpenCafeMol

OpenCafeMol is an OpenMM-based MD simulator used for CG biomolecular systems. Owing to its high performance and multiplatform OpenMM library, OpenCafeMol was developed for high-performance MD simulations, particularly for GPU-based simulations. Currently, it supports a few residue-resolution protein models and a lipid model that is compatible with the protein models.

Protein models that can be run with OpenCafeMol are C_α_-based protein models, including an atomic interaction-based CG model AICG2+ (11), an original off-lattice Go-model developed by Clementi et al. (33), and an elastic-network model (34, 35). By preparing suitable input files, several additional CG models can be simulated using OpenCafeMol. For lipids, OpenCafeMol currently implements an original version of iSoLF(25) that represents one phospholipid molecule with five particles. In addition, several standard interactions were implemented, including the Debye–Hückel electrostatic interactions, the HPS model, restraining potentials for the centers of mass of the two-particle groups, and the particle positions (26).

OpenCafeMol supports Langevin integrators with different ensembles such as NVT, NPT, and NP_γ_T ensembles, in addition to simple energy minimization. Note that the GENESIS input assumes a time unit of picoseconds, whereas the TOML input assumes a time unit of one café-time = 49 fs (see the next paragraph) (see the next paragraph for the GENESIS and TOML inputs). The recommended time steps were 0.01 (ps) with the GENESIS input, and 0.2 (café-time) with the TOML input.

OpenCafeMol was developed using the C++ API of the OpenMM Library layer (Fig. 1). It reads the input information using either TOML-style input files (defined in Mjolnir. TOML is a generic-style input data format usable for broad purpose, but not very user-friendly) or GENESIS input files (defined in GENESIS and is a user-friendly format. This is supported only partially at present). Potential energies were determined from the input information. Every potential energy term in CG models is encoded by the “class” of OpenCafeMol, which was defined by the use of the OpenMM class, as in Table I. Whenever necessary, users can add new types of potentials by creating a new class. Thus, it has a high expandability. To run the MD simulations, OpenCafeMol Simulator calls OpenMM Integrator and performs Langevin dynamics.

**Table I.**
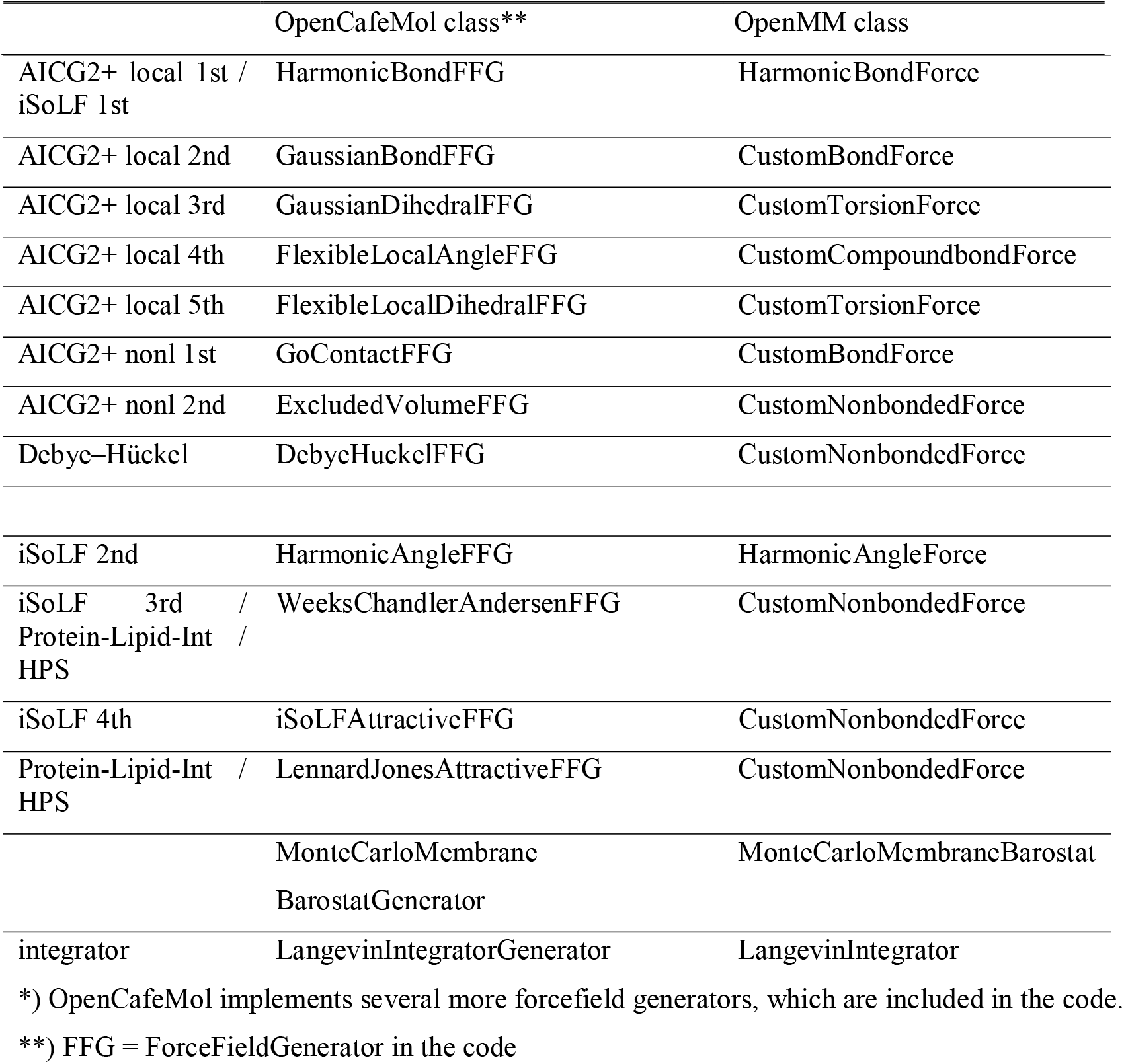
OpenCafeMol class*.

| OpenCafeMol class** | OpenMM class |
| --- | --- |

|  |  |  |
| --- | --- | --- |
| AICG2+ local 1st /<br>iSoLF 1st | HarmonicBondFFG | HarmonicBondForce |
| AICG2+ local 2nd | GaussianBondFFG | CustomBondForce |
| AICG2+ local 3rd | GaussianDihedralFFG | CustomTorsionForce |
| AICG2+ local 4th | FlexibleLocalAngleFFG | CustomCompoundbondForce |
| AICG2+ local 5th | FlexibleLocalDihedralFFG | CustomTorsionForce |
| AICG2+ nonl 1st | GoContactFFG | CustomBondForce |
| AICG2+ nonl 2nd | ExcludedVolumeFFG | CustomNonbondedForce |
| Debye-Hückel | DebyeHuckelFFG | CustomNonbondedForce |
| iSoLF 2nd | HarmonicAngleFFG | HarmonicAngleForce |
| iSoLF 3rd /<br>Protein-Lipid-Int /<br>HPS | WeeksChandlerAndersenFFG | CustomNonbondedForce |
| iSoLF 4th | iSoLFAttractiveFFG | CustomNonbondedForce |
| Protein-Lipid-Int /<br>HPS | LennardJonesAttractiveFFG | CustomNonbondedForce |
|  | MonteCarloMembrane<br>BarostatGenerator | MonteCarloMembraneBarostat |
| integrator | LangevinIntegratorGenerator | LangevinIntegrator |

\*) OpenCafeMol implements several more forcefield generators, which are included in the code.
\*\*) FFG = ForceFieldGenerator in the code

**Figure 1.**
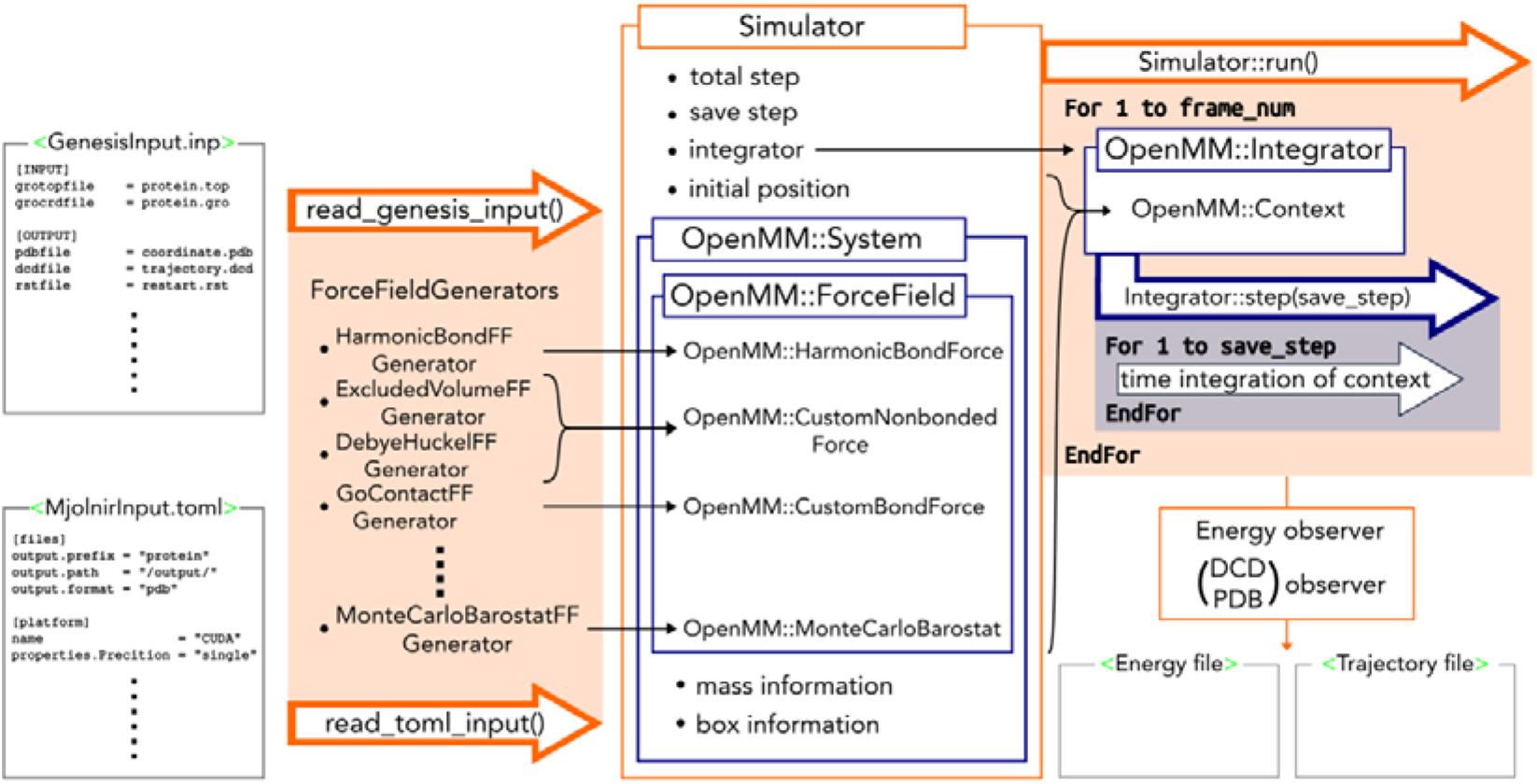
Schematic view of the OpenCafeMol software architecture.

From the user’s perspective, OpenCafeMol has two interfaces: the GENESIS and TOML (Mjolnir) interfaces. The GENESIS interface is simple, and its input format is essentially the same as that of GENESIS, which resembles the GROMACS input file format (36). Users who are unaware of the details of the force field can begin the simulation using this interface. The TOML-style input allows for more general types of potential that follow the file format of Mjolnir (https://github.com/Mjolnir-MD/Mjolnir). The TOML input file describes the detailed parameters of the force field so that users can fine-tune the force field. OpenCafeMol reads these two types of input files and translates them into the force field classes of OpenCafeMol, which generates OpenMM force fields.

### Benchmark: protein folding with AICG2+ and electrostatic interaction

First, we evaluated OpenCafeMol for the folding of a small protein, the 56-residue Src homology 3 (SH3) domain, using the AICG2+ model of proteins with Debye-Hückel electrostatic interactions (11). The AICG2+ model is a structure-based residue-resolution CG protein model that has long been used in large-scale simulations. In addition, we added electrostatic interactions between charged residues (+1 for Arg and Lys, and -1 for Asp and Glu).

As expected, we observed temperature-dependent repeated folding and unfolding transitions, which were quantified as the time series of the Q-score, a fraction of the native contacts formed (Fig. 2a). The average Q-score curve as a function of temperature around the folding–unfolding transition is very sensitive to the model and simulation setup; thus, it can be used as a stringent test for MD simulators. We calculated the average Q-scores as a function of temperature using OpenCafeMol and CafeMol from 20 independent 10^7^ MD step trajectories in each simulation (Fig. 2b). The denaturation curves from the two simulators agreed well, indicating that OpenCafeMol can accurately simulate protein folding.

**Figure 2.**
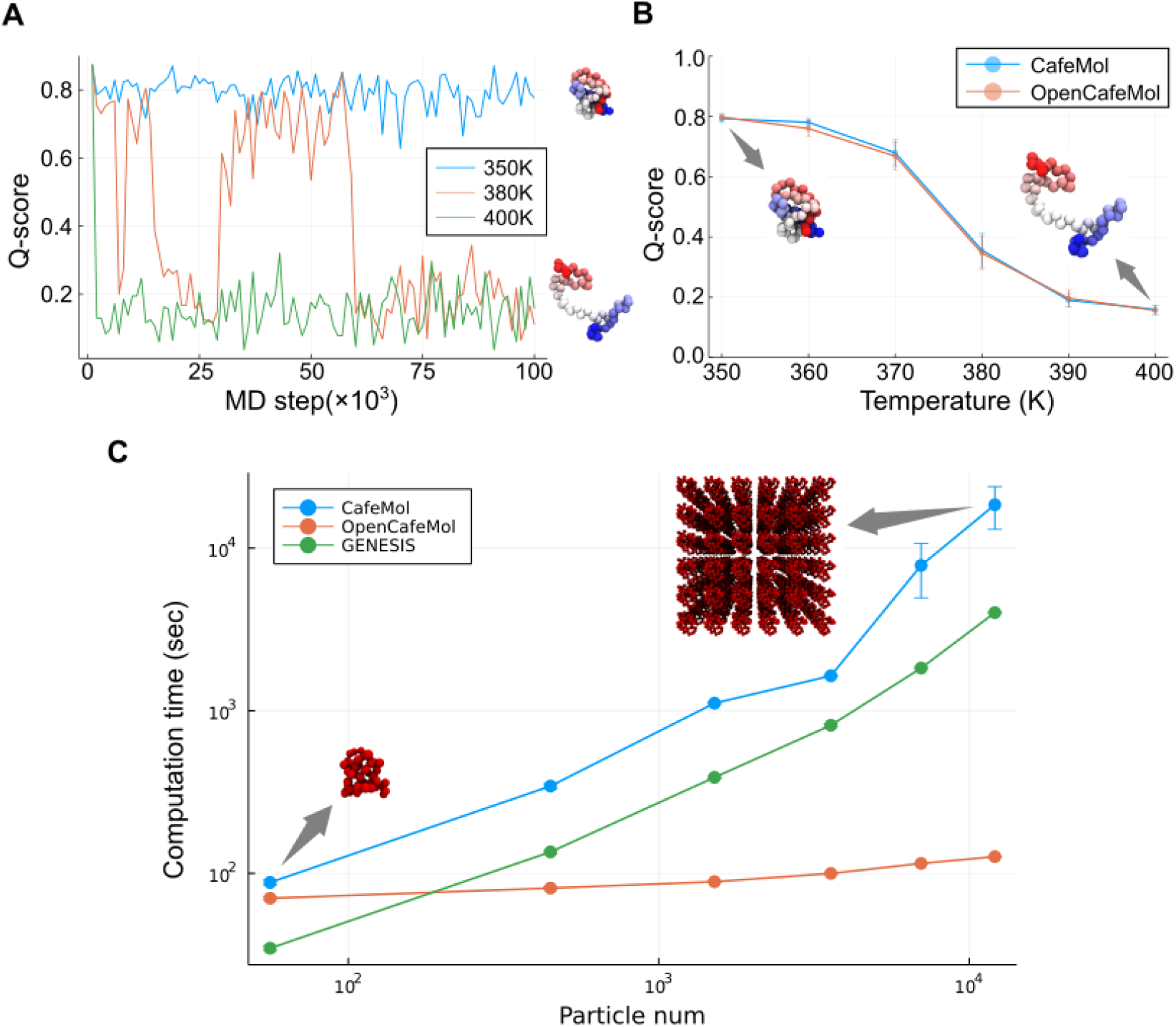
Benchmark test for folding of a small protein, the SH3 domain. (a) Representative time courses of Q-score at three temperatures. (b) The average Q-score as a function of temperature. Error bars represent the standard error. At each temperature, we ran 20 trajectories of 10^7^ MD steps. The average Q-score was obtained from data at every step in the latter half of each trajectory. (c) The benchmark of computation time for different size systems. Each data point contains 1 (56), 8 (448), 27 (1512), 64 (3584), 125 (7000), and 216 (12096) SH3 molecules (particles). The computation times were averaged over four trajectories. The error bar represents the standard error.

We then measured and compared the calculation speed of OpenCafeMol with a GPU (RTX4090 1GPU) and the previously developed simulators, CafeMol.3.2.1, and GENESIS.1.7.1, with CPUs (Intel Xeon Gold 6226 1CPU, 8 cores, MPI parallelization; It should be noted that parallelization using OpenMP results in markedly slower calculation speeds.). To examine the scalability of the simulations, we placed SH3 (PDBID:1SRL) molecules in 3D cubic grids containing 1, 8, 27, 64, 125, and 216 SH3 molecules (the number of amino acids (particles) corresponded to 56, 448, 3584, 7000, and 12906, respectively). The computation time required for 10^6^ MD steps was measured (Fig. 2c). For one SH3 domain, OpenCafeMol was only 1.26 times faster than CafeMol, whereas for the largest tested case of 216 proteins, OpenCafeMol was ∼100 times faster than CafeMol and GENESIS, which used CPUs.

### Benchmark for lipids: iSoLF model

Next, we tested OpenCafeMol for POPC lipid bilayer membrane dynamics using the CG lipid force field, iSoLF (25). The iSoLF model represents a POPC lipid with five particles: two particles for the head and three particles for the tails. In lipid membrane MD simulations, the area per lipid is sensitive to lipid dynamics, changing its mean value upon transition from the liquid-ordered phase to the liquid-disordered phase. Thus, the area per lipid molecule was used to test the simulation accuracy. We calculated the area per lipid of the POPC membrane in the temperature range of 270 K–400 K using OpenCafeMol and CafeMol (Fig. 3a). For each case, we performed 10^7^ step MD simulations and sampled structures every 10^5^ steps after the initial equilibration of 5.0×10^6^ steps. The area per lipid as a function of temperature agreed well between OpenCafeMol and CafeMol. We conclude that the iSoLF model in OpenCafeMol is accurate.

**Figure 3.**
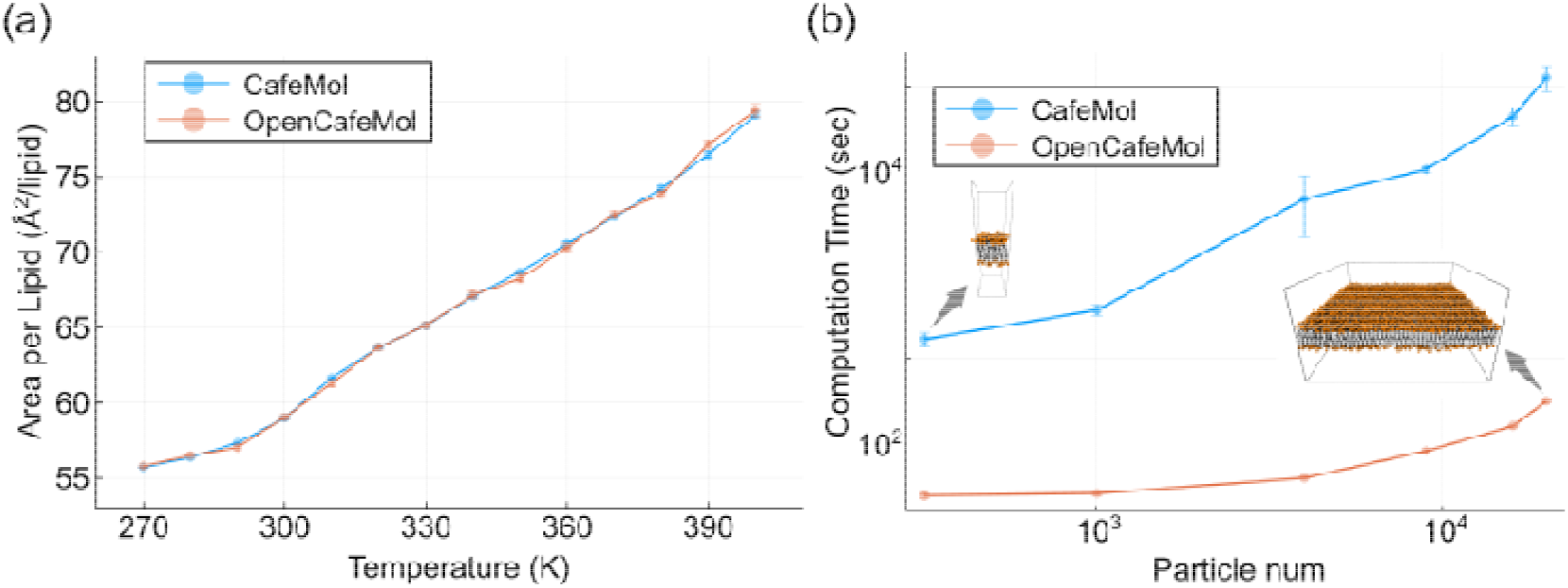
Benchmark test for the POPC lipid bilayer membrane. (a) Area per lipid as a function of temperature. For each point, the data point is the average of every 10^5^ steps after the initial step from 10^8^ MD step simulations. Error bars correspond to standard errors. (b) The benchmark result of POPC membrane simulation.

We then benchmarked the computation time of the iSoLF model using OpenCafeMol with GPUs and CafeMol with CPUs (Fig. 3b). For this purpose, we used a series of square POPC lipid bilayer membranes containing 128, 200, 800, 1800, 3200, and 5000 POPC molecules. The numbers of particles in each case were 640, 1000, 4000, 9000, 16000, and 20000. Using the same hardware as above, the calculation speed of the GPU-based OpenCafeMol was 14.1 times faster than that of the CPU-based CafeMol for the system with 128 lipids, whereas the speed-up ratio reached 239 times that of the 5,000-lipid system.

### Benchmark for protein in a lipid membrane

Third, we examined OpenCafeMol as a membrane protein, rhodopsin (PDBID:5AX0), embedded in a POPC lipid membrane using the iSoLF model and the protein-lipid interaction developed by Ugarte et al. (28). The test system comprised a unit that included one rhodopsin molecule embedded in 94 POPC molecules (Fig. 4a, left). First, we examined the distribution of rhodopsin tilt angle with respect to the membrane plane (Fig. 4ab). The overall distribution of OpenCafeMol was consistent with that of CafeMol. The distribution was slightly sharper, which can be attributed to a slight difference in the surface tension of the membrane.

**Figure 4.**
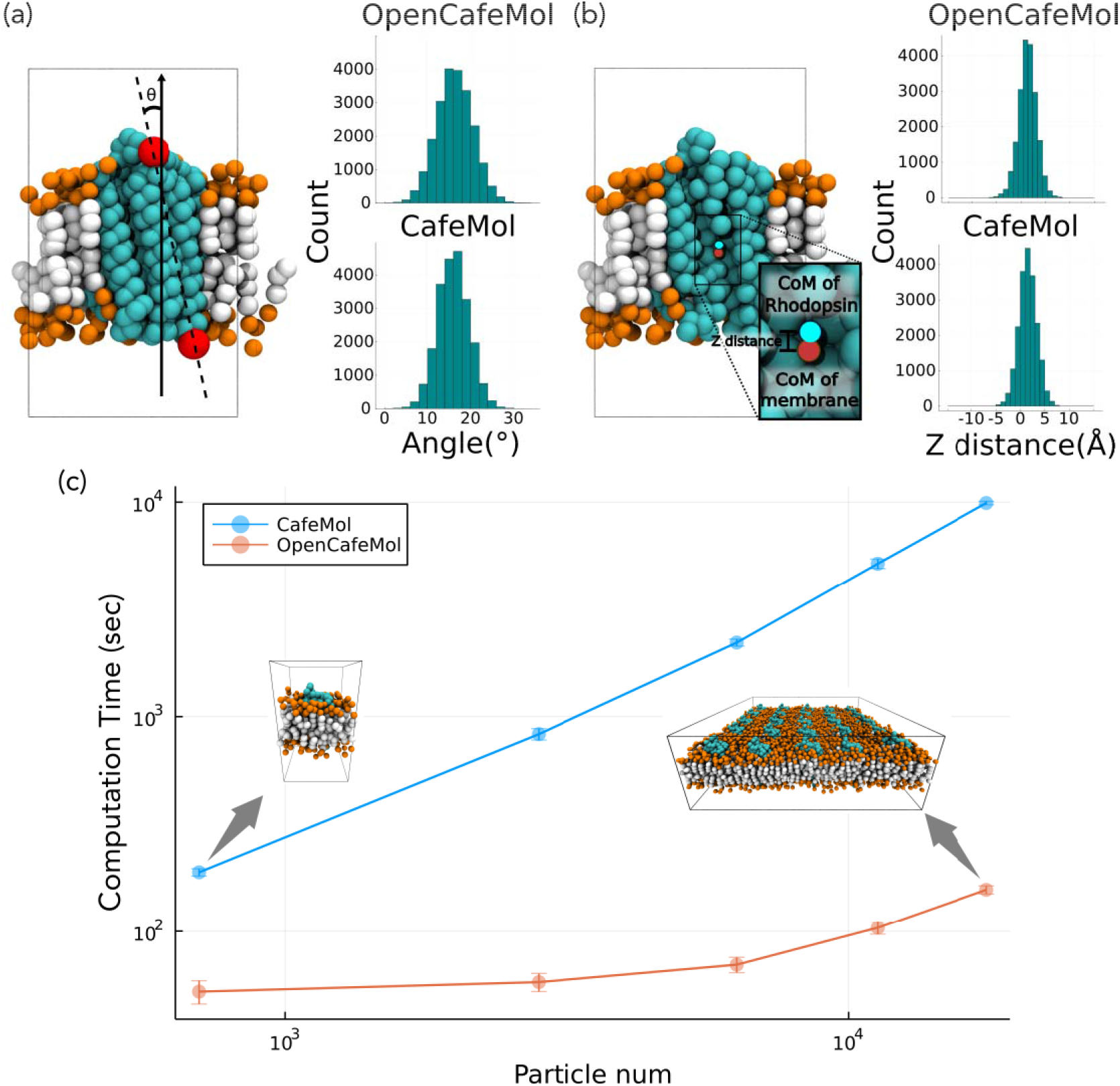
Benchmark text for a protein-lipid mixture. (a) Distribution of rhodopsin tilt angle to the normal of the membrane surface. Top: OpenCafeMol; bottom: CafeMol. (b) Distribution of rhodopsin insertion to the membrane. Top: OpenCafeMol; bottom: CafeMol. (c) Computation time of lipid-protein system simulation. Each data point corresponds to 1 (704), 4 (2816), 9 (6336), 16 (11264), and 25 (17600) units (particles).

To benchmark the computation time, we prepared a series of systems containing 1, 4, 9, 16, and 25 units. In the case with one unit, OpenCafeMol was just 3.6 times faster than CafeMol, whereas the speed-up ratio reached 63 for the 25-unit system.

### Vesicle fusion by force

Taking advantage of the high speed of OpenCafeMol, we now apply it to simulate longer dynamics of larger systems; we choose a membrane fusion of the two liposomes for this purpose. A comparison from short test runs suggests that if we had used a CPU version of CafeMol, the following simulations would have taken longer than a year. It took less than a day with OpenCafeMol with a GPU computer (RTX4090).

Each liposome is composed of 5710 POPC molecules, the diameter of which is about 50 Å (Fig. 5a). To induce membrane fusion, we set pseudo-particles at the center of each liposome and pulled them using a harmonic potential for inter-pseudo-particle distance, with no short-range repulsion between the two pseudo-particles. We illustrate a representative trajectory of the fusion process. Two vesicles made contact around the 8.0×10^5^-th MD step, which was followed by the fusion of two vesicles around the 1.6×10^6^-th MD step (Fig. 5bc). The peanut-shaped fused vesicle slowly changed shape to a near-spherical form during the subsequent 10^7^ MD steps. The homogenization of lipids took a rather long time and was completed near the end of 10^8^ MD step simulations (Fig. 5bc).

**Figure 5.**
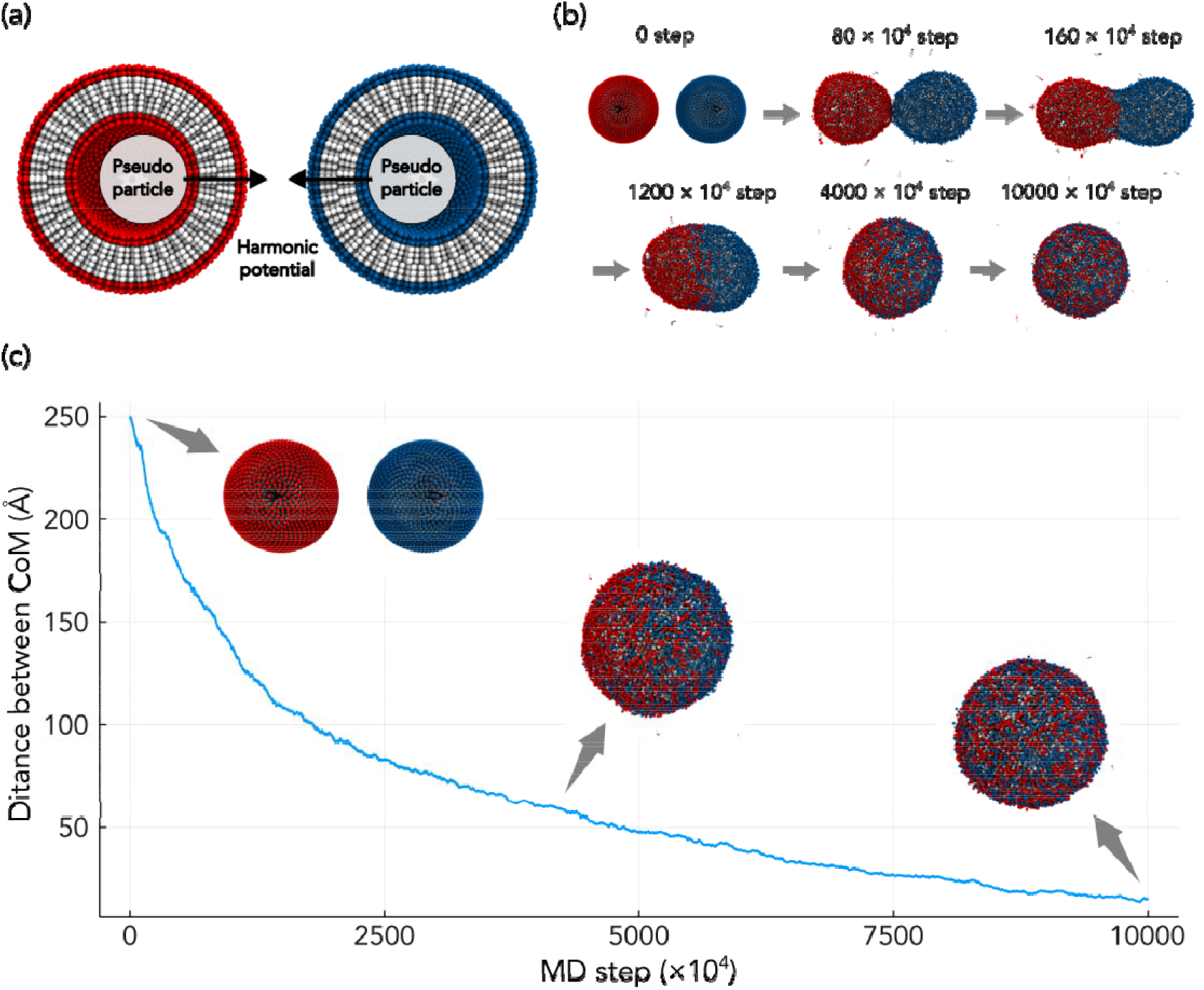
Vesicle fusion simulation. (a) The system is composed of two vesicles each with one pseudo-particle in the central cavity, used for pulling. (b) Snapshots of vesicle fusion. (c) A representative trajectory of the liposome merging process. The vertical axis is the distance between the centers of masses of lipids initially contained in two (red and blue) vesicles.

### Vesicle fusion by SNARE complex

Next, we step forward to simulate vesicle fusion in a setup closer to cellular context; vesicle fusion mediated by SNAREs (Soluble N-ethylmaleimide-sensitive factor attachment protein receptors). SNARE is a protein family that mediate membrane fusion and is conserved across *Eukaryote* (37). Among them, a particularly well studied system is a SNARE in neuronal exocytosis which consists of Syntaxin-1, SNAP-25, and vesicle-associated membrane protein 2 (VAMP2, a member of Synaptobrevins). These three subunits form a SNARE complex with a four-α-helix bundle structure; SNAP-25 contains two *α* -helix and each of VAMP-2 and Syntaxin-1 has one α-helix followed by a short linker and a transmembrane helix anchored to membrane to be fused. Before the fusion, they are assumed to take a *trans*-SNARE complex in which a transmembrane helix of Syntaxin-1 and that in VAMP-2 are anchored to the plasma membrane of the pre-synapse and the membrane of a synaptic vesicle, respectively. During the fusion, the complex is supposed to zip into the fully assembled *cis*-SNARE complex (29, 38–41). In *cis*-SNARE complex, the short linker between the α-helix in the bundle and the transmembrane region forms a continuous *α* -helix. There is a model in which this helix formation considered to exert force to induce membrane merge. Although there are some coarse-grained and all-atom MD simulation studies which support this model, the details still remain unclear (42–44). Notably, Risselada et al used a MARTINI coarse-grained model to successfully induce vesicle fusion by SNARE, providing detailed structural views. However, the use of an elastic network model for SNARE restrained to the cis-SNARE configuration might have enforced the fusion (42).

Now, using OpenCafeMol we simulate vesicle fusion process driven by a SNARE complex. For the two vesicles, we used the same configuration as that in the previous section. For the SNARE complex, we prepared a *trans*-SNARE complex structure by pulling transmembrane helices of Synaptobrevin (VAMP2) and Syntaxin-1A to the two opposing vesicles. Each of two transmembrane helices is inserted into each of two vesicles, thus linking them (Fig. 6a, 0 step). Importantly, the use of AICG2+ model for SNARE enables a linker of SNARE to locally unfold making a *trans*-SNARE configuration stable. With this initial configuration, we performed two independent runs of 10^8^ MD steps at each of temperatures, 303K and 363K.

**Figure 6.**
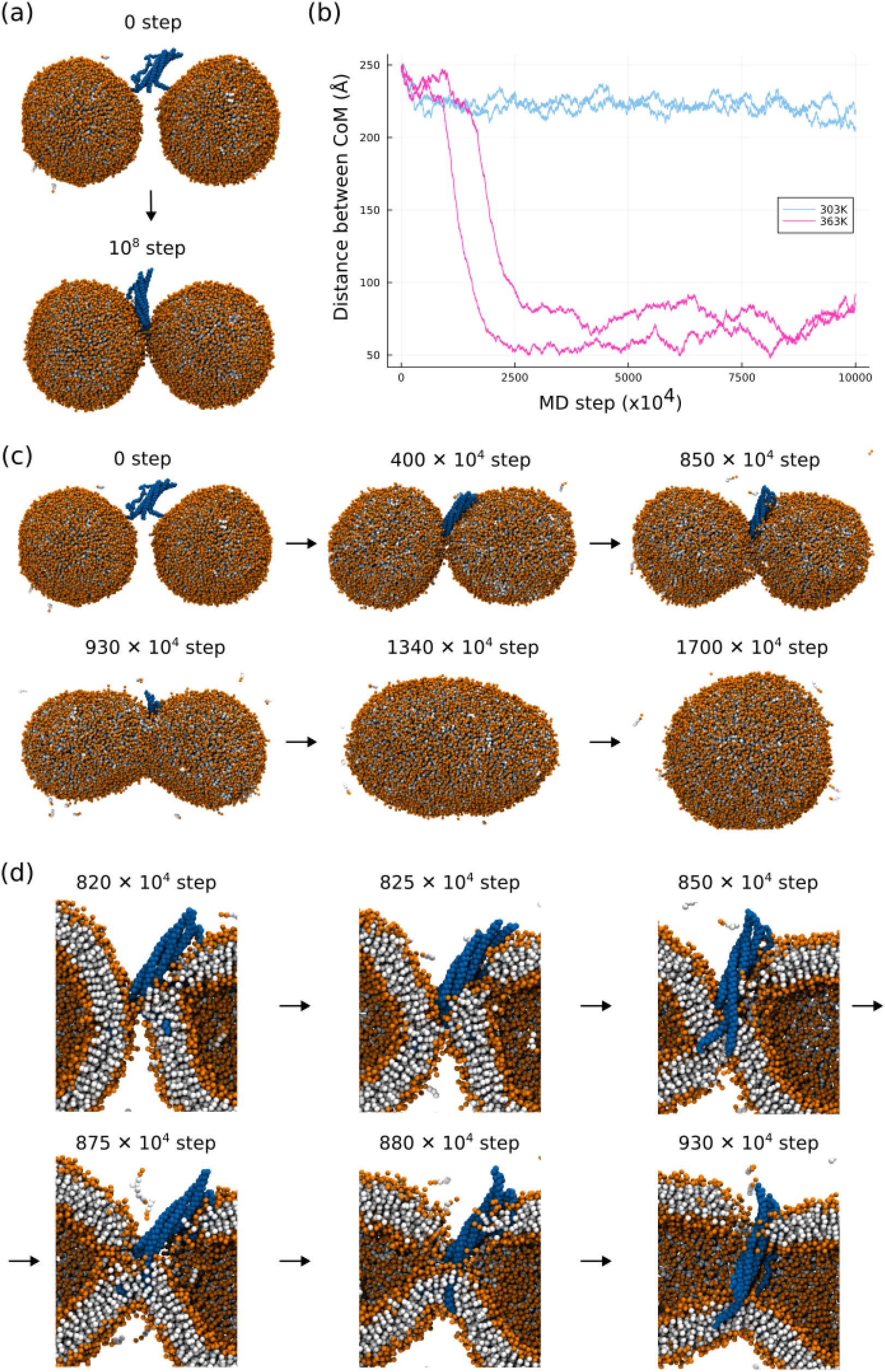
Vesicle fusion driven by a SNARE complex. (a) Representative snapshots in the simulation at 303K. (b) The time course of the distance between centers of mass of lipids that belong to the left and right vesicles in the initial configuration. Results at 303K and 363K are shown in cyan and magenta, respectively. (c) Representative snapshots in the simulation at 363K. (d) Close-up and cross-section views of the fusion junction during the fusion process at 363K. In (a), (c), and (d), lipid head, lipid tail, and SNARE complex are colored by orange, white, and blue, respectively.

Figure 6a shows the initial and final snapshots in a run at 303K. The two initially-separated vesicles quickly reached to the docked state and stayed in that state until the end of simulation. We did not observe vesicle fusion event in either trajectory. We plot time courses of the distance between centers of mass of two groups of lipids; those in the left and right vesicles at the beginning (Fig. 6b). The distance decreased by ∼20 □ at the very beginning, and then fluctuated around ∼220 Å until the end of the simulation. In the docked state, while the helix bundle of the SNARE complex is completely formed, the linkers between SNARE complex and the transmembrane helices of Synaptobrevin and Syntaxin-1A were disordered.

On the other hands, we observed vesicle fusion in the simulations at 363K. Figure 6c shows representative snapshots in a run at 363K. Two vesicles located in a separated position in initial configuration came close each other and they docked each other around 4000 × 10^4^-th step. This docked state lasted until ∼820 × 10^4^-th MD step, around when lipid tail splaying process was observed (Fig. 6d). Then, a fusion pore was formed at around 880 × 10^4^-th MD step (Fig. 6d). During this process, we did not observe the so-called hemi-fusion; Either the hemi-fusion did not occur or it was so transient that we did not observe. Before the fusion (pre-fusion), the *trans*-SNARE maintained the helix bundle in the core of the complex, but exhibited marked disorder in the linkers between the core the transmembrane helices (Fig. S1 ab). After the fusion (post-fusion), these linkers got ordered to form longer helices in the *cis*-SNARE configuration (Fig. S1b, right). Thus, local folding, or disorder-to-order transition in the linker regions can provide the driving force for the vesicle fusion. In the other run at 363K, we observed qualitatively the same sequence of events in the vesicle fusion as those described above.

## DISCUSSION AND CONCLUSIONS

In this study, we developed a fast and flexible MD software, OpenCafeMol, for residue-resolution protein and lipid models, which exhibited high performance on GPU machines. We validated OpenCafeMol for the folding of a small protein using the AICG2+ model, lipid membrane dynamics using the iSoLF model, and membrane protein structure. Benchmark tests of the computation times showed that OpenCafeMol with one GPU (RTX4090) for proteins and lipid membranes was approximately 100 and 240 times faster than the corresponding simulations with a typical CPU (Intel Xeon Gold 6226, 1CPU, 8 cores), respectively.

As new software, there is much room for extension in OpenCafeMol. A DNA model that is compatible with the residue-resolution protein model is highly desirable. In a separate article, we are reporting the implementation of the 3SPN.2C model of DNA; the 3SPN.2C model approximates one nucleotide as three particles, each representing phosphate, sugar, and nucleobase, and has broadly been used in the community (45). In addition, we can simulate protein-DNA complexes with OpenCafeMol on a GPU, which is orders of magnitude faster than CafeMol on a CPU. For proteins, we implemented a structure-based model, such as AICG2+, for protein dynamics simulations in one state. Often, to simulate transitions between two or more states, multiple-basin models are useful (46) but have not been implemented. For lipid models, we implemented only the first version of iSoLF; however, a much more extended version has been developed (47) that needs to be implemented. For time integration and structural sampling, we implemented only the Langevin dynamics at a constant temperature. Therefore, the implementation of advanced sampling methods is highly desirable. Currently, the GENESIS input files are only partially supported. We are working to support the GENESIS input files.

While the simulations of vesicle fusion by a SNARE complex gave apparently promising results, we need some caution on the interpretation and need to discuss the limitations.

First, we observed vesicle fusion events only at a high temperature, but not at a physiological temperature. One simple explanation is that a high temperature accelerated the fusion event so that the fusion occurred in the simulation time window of 10^8^ MD steps at 363K, but the simulation at 303K was too short to observe the fusion. If we could simulate much longer time at 303K, we would have possibly observed the fusion. However, a high level of fluctuations of lipids may have destabilized the vesicle markedly and induced break-up of vesicles possibly as an artifact. We confirmed that, without the SNARE complex, the direct 10^8^ MD steps-simulations at 363K does not spontaneously induce the vesicle fusion. Thus, it is the effect of SNARE that drove the fusion. However, non-physiological effect at an elevated temperature cannot be excluded.

Second, Fig. 6d shows intriguing sequence of events, which resulted in the SNARE complex inside, but not the outside, the fused vesicle. This inside-configuration of the SNARE complex after the fusion is rather unexpected. In the context of fusion of synaptic vesicle to the pre-synapse plasma membrane, the current inside-configuration of the SNARE is mapped to the configuration where the SNARE complex is located outside of the cell. This final configuration obtained from the current simulation may be different from the standard view. Notably, during the pore formation process, Fig. 6d suggests that the two transmembrane helices remain in the bottom side of the lipid bilayer (e.g. see the snapshot at 880 x 10^4^-th MD step), which resulted in the inside-configuration of the SNARE complex.

At a physiological condition, more than one SNARE complex may be necessary to achieve the vesicle fusion. We started to examine this possibility by putting two *trans*-SNARE complexes in the initial configuration. Preliminary simulations resulted in break-up of vesicles to reach one flat lipid bilayer. Our vesicles may be too small to accommodate more than one SNARE complexes. More comprehensive surveys are necessary.

While there are some possible artifacts and the limitations, the current study opens up new avenue to explore the molecular mechanisms of protein-mediated vesicle fusion process via MD simulations.

## AUTHOR CONTRIBUTIONS

Y.M. and S.T. conceived the project, and Y.M. and T.N. developed the code. Y.M. performed the simulations, analyzed the data, and assembled the figures; Y.M. and S.T. discussed the results and wrote the manuscript.

## ACKNOWLEDGMENTS

We would like to thank Masataka Yamauchi for helpful discussions. We appreciate Cheng Tan, Jaewoon Jung, and Yuji Sugita for their advice and discussions on the GENESIS implementation. This work was supported by the JSPS KAKENHI grants 20H05934 (S.T.), 21H02441 (S.T.), and 24K01991 (S.H.) by the MEXT grant JPMXP1020230119 as “Program for Promoting Researches on the Supercomputer Fugaku” (S.T.).

