## Supplementary material for "OpenCafeMol: A coarse-grained biomolecular simulator on GPU with its application to vesicle fusion": Fig. S1

**SUPPLEMENTΑL INFORMATION**

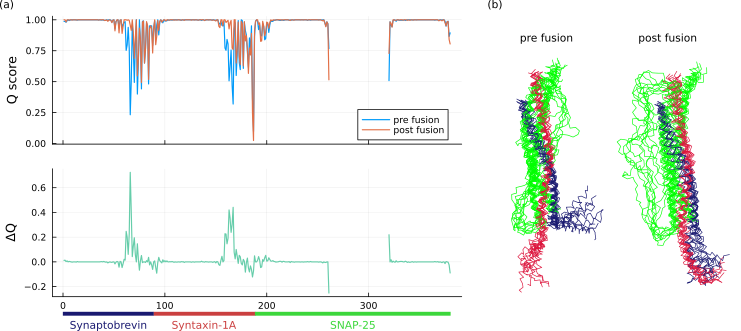

Figure S1. (a) The Q score and ΔQ score (Q_post_ – Q_pre_) between post-fusion state and pre-fusion state in 363K simulation (The Q score is defined as the fraction of contacts formed in the simulated structure to the contacts found in the reference native structure). The pre-fusion state is defined by the period from 4.0 × 10^6^ -th MD step to 8.0 × 10^6^ -th MD step and the post-fusion state is defined by the period from 9.6 ×10^7^ -th MD step to 1.0 × 10^8^ -th MD step. SNAP-25 have an IDR region in the middle of sequence, where Q score is not defined. Synaptobrevin and Syntaxin-1A have more disordered region in the pre-fusion state, compared to the post-fusion state. (b) Superposed structures of pre- and post-fusion states. In the pre-fusion state, the linkers between the helix bundle and transmembrane helices exhibit disorder. On the other hands, in the post-fusion state, these linkers fold into helices as in the *cis*-SNARE configuration. These area correspond to the ΔQ peak in the lower figure of (a).
